# Effect of carrier droplet size and shape at different viral loads on virus stability during environmental drying

**DOI:** 10.1101/2025.03.27.645702

**Authors:** Sesan Nayak, Y.S. Mayya, Kiran Kondabagil, Mahesh S. Tirumkudulu

## Abstract

In the aftermath of the COVID-19 pandemic, airborne transmission has been identified as a significant factor in disease spread. However, there have been very few direct comparisons of virus viability in airborne droplets versus those deposited on surfaces or fomites. This study compares the viability of the enveloped Phi6 virus and two non-enveloped viruses (T4 and MS2) in droplets on hydrophobic and hydrophilic surfaces at 25°C and 45%–55% relative humidity. The former results in spherical droplets similar to airborne droplets, while the latter pertains to spreading droplets comparable to a fomite state. Our research highlights the influence of various physical factors of the carrier droplet—such as its shape and size—as well as the type and concentration of the virus on its viability during the drying process.

We found that, at a fixed volume, virus viability decreased with droplet size in spherical droplets, while high initial viral concentrations (∼ 10^5^ pfu/µL) improved survival on fomites. This suggests that fomites from individuals with high viral loads pose a greater risk of infection. Overall, droplets of a specific volume are more viable in the air than on surfaces. Smaller airborne droplets may have decreased viability but can linger longer and penetrate deeper into the respiratory tract. When viral loads are high, its comparable persistence in both spherical and flat droplets increases the risk of fomite transmission.

**IMPORTANCE:** Understanding the roles of airborne and fomite transmission in respiratory diseases is essential for planning effective containment measures and resource allocation. Among the various factors that influence the spread of infection, the survival of viruses in drying droplets plays a critical role in disease transmission. In our study, we compare the viability of viruses in spherical droplets, which represent airborne transmission, and flat droplets, which represent fomite transmission. Our results indicate that virus viability is consistently higher in spherical droplets than in flat droplets when dried under similar conditions. However, for high virus loads, the difference in virus survival is significantly lower. Therefore, during an outbreak, it is important to monitor both the number of new infections and the viral load of infected individuals, as those with higher viral loads are more likely to spread the infection through both modes of transmission.

## INTRODUCTION

The frequent global outbreaks of respiratory disease present a significant concern across the world. COVID-19 pandemic caused considerable disruption from 2019 to 2021 before the coronavirus vaccine became available, and it continues to affect populations, although fatalities are now more manageable. As of early 2025, there are increasing concerns regarding an HMPV outbreak, which is currently confined to China.

Airborne transmission is primarily responsible for the spread of the coronavirus, as infectious viruses shed from a source reach susceptible hosts through respiratory (i.e. saliva or mucus) droplets and aerosols (1). Additionally, virus-laden droplets that land on surfaces contribute to indirect transmission, known as fomite transmission. Among various physico-bio-environmental factors that influence the spread of infection, the persistence or survival of viruses in droplets that dry in the ambient environment plays a critical role in transferring the infection and causing disease.

Reliable data pertaining to virus viability during environmental drying is still inadequate. To evaluate the potential impact of an outbreak, disease modelers and policymakers often resort to conservative estimates of viral load survival rates using arbitrary infectivity parameters. Thus, it is crucial to further investigate this issue to achieve a deeper understanding of outbreak occurrences and to formulate effective strategies to combat them.

In the early days of the COVID-19 pandemic, there was a significant focus on surface disinfection and hand sanitization as preventive measures against transmission. This emphasis stemmed from the prevalent belief that contact-based and surface-based transmission, commonly known as fomite transmission, was more critical than airborne transmission. However, this notion was later challenged on grounds that small droplets exhaled by an infected person can travel significant distances in air and carry their viral load (2,3). Subsequent studies from various outbreak investigations confirmed that airborne transmission was, in fact, the dominant mode of COVID-19 spread (4,5). While larger droplets, which make up the majority of expelled respiratory fluid by volume, can contaminate surfaces and contribute to fomite transmission, there is a lot of variability in the literature regarding how long viruses can survive on surfaces and maintain their infectivity (6,7,8,9).

Numerous studies and analyses, both experimental and theoretical (10,11), along with several well-cited reviews (12,23), have strongly supported the view that airborne transmission predominates. However, these studies often focus on outbreak investigations and generally lack direct comparisons between airborne and fomite infectivity.

It is generally accepted that the survival of viruses is significantly influenced by their microenvironment, which is shaped by the physiochemical characteristics of the drying matrix (e.g., droplet composition) as well as the drying conditions (e.g., temperature and relative humidity). Salts and proteins are major non-volatile components of saliva, and studies have reported that these components have varying effects on viruses depending on the strain and environmental conditions (14, 15, 16). Additionally, research has indicated that a high initial viral load or concentration in drying droplets can protect the resident viruses (17,18). However, the differences in the effects of initial viral load on the viability of airborne versus fomite viruses have not been adequately explored.

The carrying capacity of droplets, which depends on their size, is considered a significant factor in their infectivity (19). Additionally, droplet size plays a crucial role in determining the drying rate. As a droplet shrinks to its final stable residue particle, the drying stress experienced by viruses can also affect their survival. Thus, droplet size can impact the overall transmissibility of viruses in multiple ways.

First, there are size-dependent Poisson fluctuations concerning the likelihood of virus incorporation into the droplet (20). This is further influenced by the shielding effect of the liquid on the virus’s infectivity. Zuo et al. found that droplets larger than the size of the viruses (300-450 nm) were more infectious (21). In other words, there was a higher virus recovery, compared to droplets that were closer in size to the viruses (100-200 nm). The authors attribute this phenomenon to the shielding effects provided by the larger droplets.

However, it’s important to note that in smaller airborne droplets, the shielding effect may be limited, as these droplets typically shrink to solid residue particles that are nearly one-fifth of their original size (22). In another study, Alonso et al. reported that viral viability was higher in larger carriers for three different porcine viruses; specifically, the influenza A virus, the porcine reproductive and respiratory syndrome virus, and the porcine epidemic diarrhea virus, each with various transmission routes (23).

Quantifying this survivability under various parameters of the virus and its carrier droplet, can offer valuable insights into transmission risks in different scenarios. In this study, we focused on larger settled droplets instead of the smaller airborne droplets (which are generally less than 100 µm) because they are easier to handle and allow for better recovery of residual viruses. While experiments conducted in controlled environmental chambers can generate relevant-sized micro-droplets or aerosols, the recovery of viable viruses from these micro-droplets is often complicated and results in significant loss of viability (24,25,26). The recovery process must account for losses caused by the techniques or equipment used (such as impingers, impactors, and filters) and the unaccounted settling of aerosols. These uncertainties make viability studies that involve airborne droplets directly quite unreliable. In light of these challenges, several previous studies have opted to use sessile droplet drying on surfaces to quantify virus decay under different conditions. This approach has been favored for its reliability and convenience (18,27,28,29).

Here, we employed the sessile drop drying approach to evaluate the impact of various parameters on virus viability losses. The factors investigated included virus type (enveloped vs. non-enveloped), droplet composition (salt-rich vs. protein-rich), droplet shape & size, and initial virus concentration. Additionally, we explored the differences in viral viability between airborne droplets and those on surfaces (fomites) under similar drying conditions, using a simple experimental setup. To understand the effect of carrier droplet size on the viability of resident viruses, we dried droplets of different sizes while maintaining consistent conditions.

## MATERIALS AND METHODS

### Types of viruses used

Viruses generally consist of genetic material (either DNA or RNA) encased in a protective protein coat known as the capsid. Some viruses also possess an additional lipid layer called a lipid envelope. This distinction between having or lacking a lipid envelope classifies viruses into two categories: enveloped and non-enveloped viruses. Respiratory viruses, such as influenza, coronaviruses, and RSV, are typically enveloped viruses.

In our study, we utilized a surrogate for COVID-19, known as Phi6 phage, which is an enveloped RNA virus. Phi6 is commonly employed as a surrogate due to its outer lipid envelope and its safety profile for humans (18, 30). Additionally, we experimented with T4 and MS2 phages as models for non-enveloped viruses to evaluate the impact of the lipid envelope on viral survival during environmental drying. A comparison of the three viruses is presented in Table 1, along with a schematic representation in Figure 1.

**Table 1:** Comparison of the phages across their different attributes.

| Feature | T4 | MS2 | Phi6 |
| --- | --- | --- | --- |
| Genome | Double-stranded DNA (dsDNA). ~169 kb | Single-stranded DNA (ssDNA). 3.569 kb | Double-stranded RNA (dsRNA); segmented-3 segments, 13.5 kb |
| Morphology | Icosahedral head, tail fibers | Small, icosahedral | Enveloped, icosahedral |
| Size | 200 nm long x 90 nm wide | ~23-27 nm in diameter | ~80 nm in diameter |
| Host | *Escherichia coli* B40 | *Escherichia coli* C3000 | *Pseudomonas syringae* |

**Figure 1.**
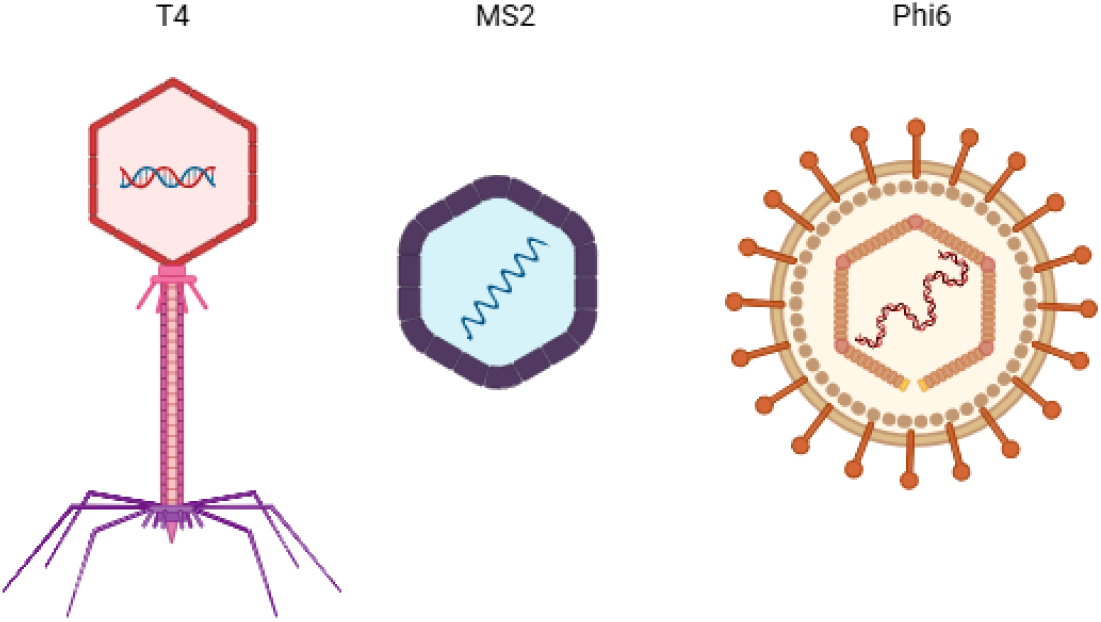
Schematic 2D images of T4, MS2 and Phi6 phage (made through Biorender.com)

### Droplet composition

To study the impact of droplet composition on resident viruses during the drying process, we used SM buffer and artificial saliva. These two solutions represent salt-rich and protein-rich media, respectively (31). The SM buffer contains sodium and magnesium salts (composed of 100 mM sodium chloride, 10 mM magnesium sulfate, 50 mM Tris-HCl at pH 7.5, and 0.01% (w/v) gelatin), while the protein content of the artificial saliva is derived from mucin (NaHCO3 5.2 g/L, NaCl 0.88 g/L, K2HPO4 1.36 g/L, KCL 0.48 g/L, α-amylase 2000 units/L, porcine gastric mucin 2 g/L) (32).

### Droplet shapes

Teflon is a hydrophobic material with a contact angle (CA) greater than 100°, causing water-based droplets to retain a spherical shape when deposited on its surface. In contrast, hydrophilic materials like glass, which have a contact angle of less than 40°, allow droplets to spread out (see Figure 1a). The hydrophobic characteristics of Teflon enable the formation of spherical droplets, which can serve as surrogates for airborne droplets due to their shape. Thus, comparing virus viability on Teflon and glass provides a useful proxy for assessing the infectivity of airborne and fomite viruses.

Spherical and flat droplets, which serve as proxies for airborne particles and fomites respectively, were created by placing equal volumes of droplets on Teflon and glass (CA approx. 105°-110° and 30°-40°) slides (see Figure 2a). It is important that the surfaces are clean and free of any visible debris, as such contaminants that can affect the droplets’ contact angle.

**Figure 2.**
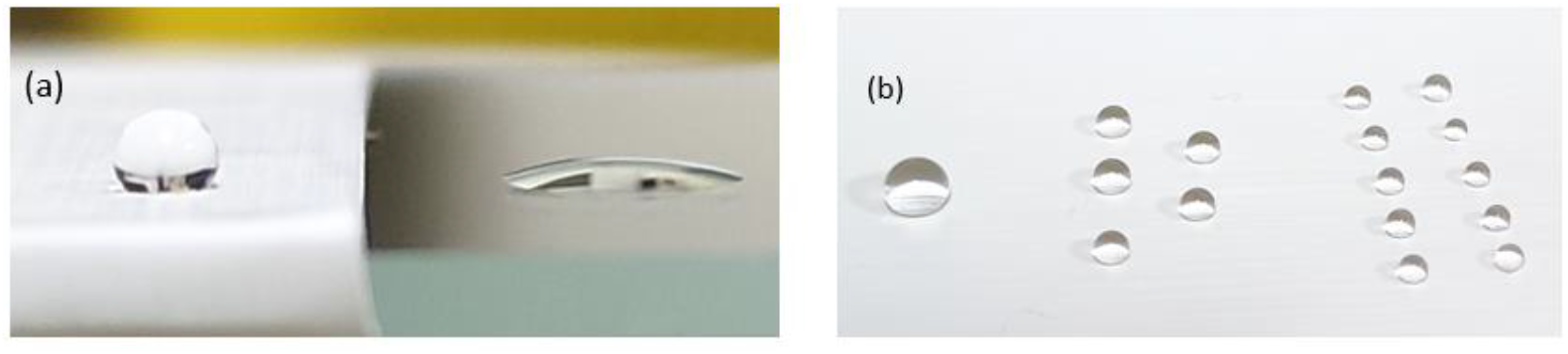
(a) 2 µL SM buffer droplet on Teflon (left), glass (right), (b) AS droplet of 10 µL (left), 5×2 µL (middle), 10×1 µL (right), on Teflon. Photos taken by smartphone camera. Droplet drying on Teflon represents conditions akin to those experienced during airborne transmission while drying of drops on glass substrate represent conditions relevant to fomite transmission

### Initial viral concentration

Stocks of Phi6 with two viral concentrations (10^3^ and 10^5^ pfu/µL) were prepared from the main stock solution through serial dilution in both SM buffer and artificial saliva. Equal volumes of these Phi6 stocks were dried on Teflon and glass surfaces under the same drying conditions and then retrieved after complete drying.

### Varying droplet size

When comparing virus viability across different droplet sizes, it is important to keep the initial cumulative droplet volume consistent. This ensures that the starting virus count among batches of droplets of varying sizes remains similar. In this study, Phi6 carrying AS droplets of different diameters but with a fixed cumulative volume were used: one drop of 10 µL, five drops of 2 µL each, and ten drops of 1 µL each. These droplets were dried simultaneously on Teflon (Figure 2b) and retrieved after approximately 60 minutes, which was sufficient time for the droplets to dry completely. The dried residues were subsequently re-suspended in SM buffer by pipetting.

### Host bacteria culture and plaque assay

The double-agar layer technique was employed for the propagation of Phi6. First, a single colony of P. syringae (PS) on an agar plate was inoculated into 5 mL of tryptone broth and incubated overnight in a shaker at 25°C and 130 RPM. Afterwards, 300 µL of the overnight culture was transferred to another 5 mL of tryptone broth and incubated under the same conditions for 8 hours to achieve the necessary concentration of host bacteria for Phi6 propagation.

Next, 300 µL of this PS culture containing approximately 10^6^ cells was mixed with 10 µL of Phi6 stock (with a concentration of 10^5^ PFU/µL) in a glass tube to achieve a multiplicity of infection (MOI) of roughly 1. This mixture was then incubated for 10 minutes at 25°C to facilitate phage infection.

Following this, 3 mL of top agar (0.7% w/v agar in tryptone) was added to the phage-bacteria mixture and thoroughly mixed by shaking. The mixture was then plated on an LB agar plate and incubated overnight at 25°C. For virus enrichment using the plate method, 3 mL of SM buffer was added to each plate. The plates were kept on a rocker at 4°C for 4 hours.

The SM buffer containing the Phi6 virion particles was then collected, filtered, and centrifuged at 20000 x g for 30 minutes at 4°C. The phage concentration in the collected supernatant was estimated using plaque assay with standard dilution methods. A similar protocol was used for the propagation of T4 and MS2 phages with their respective hosts, E. coli B40 and E. coli 3000, with the only difference being the incubation temperature of 37°C.

### Virus retrieval and viability assessment

The procedure for all virus drying experiments is as follows: A specific volume of bacterial viruses (such as T4, Phi6, and MS2) in either SM buffer or artificial saliva was deposited onto the chosen substrates, which included Teflon and glass. The large contact angle of the droplet on Teflon simulates the drying of a suspended droplet during airborne transmission, while the lower contact angle on the glass substrate represents conditions typical of fomite transmission (see Figure 1a).

The viral droplets were allowed to dry under ambient conditions, at around 25°C and 45-60% relative humidity. After a drying duration of approximately one hour, the dried residues were re-suspended in SM buffer, using a volume ten times greater than that of the original droplet, which was achieved through repeated pipetting. The recovered viruses in the SM buffer were then subjected to a standard plaque assay to quantify their viability. Each experiment was repeated at least four times.

### Data analysis

All pfu counts for the source sample were calculated using the following formula:

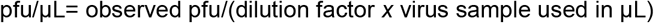

The virus pfu counts were expressed as their logarithmic values. To assess inactivation, the log-pfu counts of the experimental lot were divided by those of the control samples. In cases where no pfus were observed, a log value of 0.01 was used for calculation convenience. All data analysis was performed using Microsoft Excel, and plotting was conducted with OriginLab software.

## RESULTS AND DISCUSSIONS

Effect of droplet composition (salt vs protein) on the viability of different Virus types (enveloped and non-enveloped)

Figure 3 illustrates the decline in viability of three virus strains when suspended in SM buffer and artificial saliva (AS). Among the tested strains, the double-stranded DNA phage T4 was the most sensitive to drying, showing a decrease in the count of three to four orders of magnitude (from approximately 10^6 to about 10^3 and 10^2 in SM and AS, respectively). In contrast, the RNA viruses Phi6 and MS2 exhibited a smaller decline, roughly one order of magnitude.

**Figure 3.**
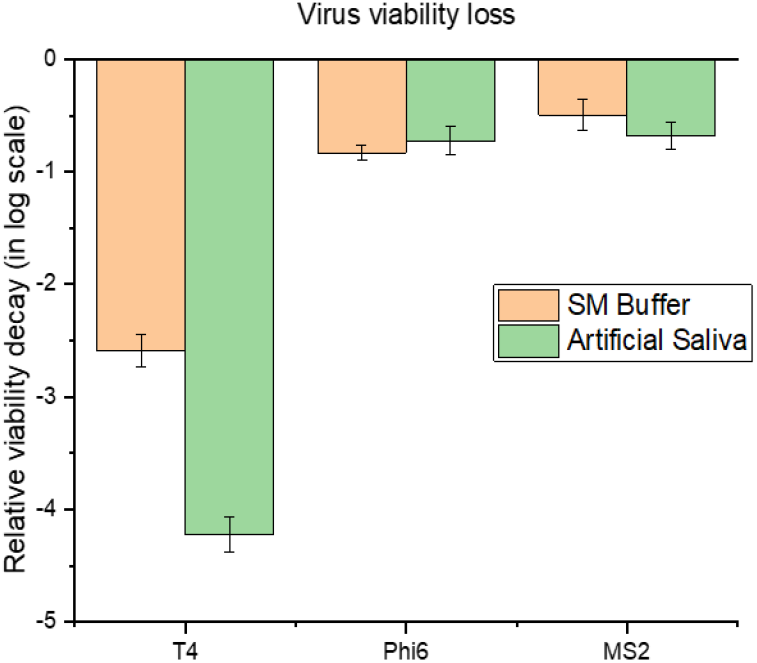
Droplet composition dependency of virus viability (all dried on Teflon)

Generally, enveloped viruses are known to be more sensitive to drying (16) because their phospholipid envelopes are affected by changes in pH and osmotic stress in drying droplets. In this study, we observed that MS2 demonstrated slightly greater persistence than Phi6. However, the heightened vulnerability of the T4 phage may be attributed to its structure. The head and tail complex of coliphages like T4 is essential for host infection, and it can sustain damage during the drying process (33). Additionally, the high DNA packaging stress in coliphages may contribute to their susceptibility to capsid bursting under stressful conditions (34).

The superior persistence of T4 and MS2 phages in SM buffer compared to AS (Figure 2) is consistent with previous studies that indicate non-enveloped viruses perform better in salt-rich environments than in salt-deficient ones (35,36). It has been suggested that salt ions stabilize non-enveloped viruses during drying by preventing structural rearrangements of their capsids. For enveloped viruses, research indicates that proteins support or enhance virus viability during drying (16,37). In contrast, salt ions are thought to interact with the lipid layer, leading to structural and mechanical changes (38,39). In our study, the viability of Phi6 was similar in both droplet compositions, with a slight increase in AS. These findings align with existing literature, providing a measure of validation for our experimental procedure.

### Virus viability on hydrophobic and hydrophilic surfaces: effect of droplet Shape

The experiments conducted on the viability of viruses retrieved from dried droplets on Teflon and glass surfaces provide significant insights. Teflon, being hydrophobic, preserved the spherical shape of the droplet due to its high contact angle, while on glass, the droplets spread out because of the lower contact angle (see Figure 1a). Under ambient conditions, 2 µL droplets from the same virus stock evaporated more quickly on the glass surface (approximately 15 minutes) compared to the Teflon surface (around 50-60 minutes). Both dried residues were successfully recovered after roughly 60 minutes of drying initiation. We examined viability loss on Teflon and glass for all three viruses, testing Phi6’s viability loss at two initial concentrations: approximately 10^5^ pfu/µL (high) and 10^3^ pfu/µL (low). The drying conditions remained consistent for both viral loads, including indoor environments, drying time, and surface type.

As discussed previously, T4 exhibited high sensitivity to drying. When dried on Teflon, T4 phages were somewhat viable; however, on glass, the T4 phage viability was negligible (see Figure 4a). In contrast, MS2 demonstrated greater tolerance to drying, showing better viral persistence on glass, although its viability was still lower than that of the viruses recovered from Teflon (see Figure 4b). A similar trend was observed for Phi6 at both viral loads (10^5 and 10^3 pfu/µL), with Teflon showing higher virus viability than glass (see Figures 4c and 4d).

**Figure 4.**
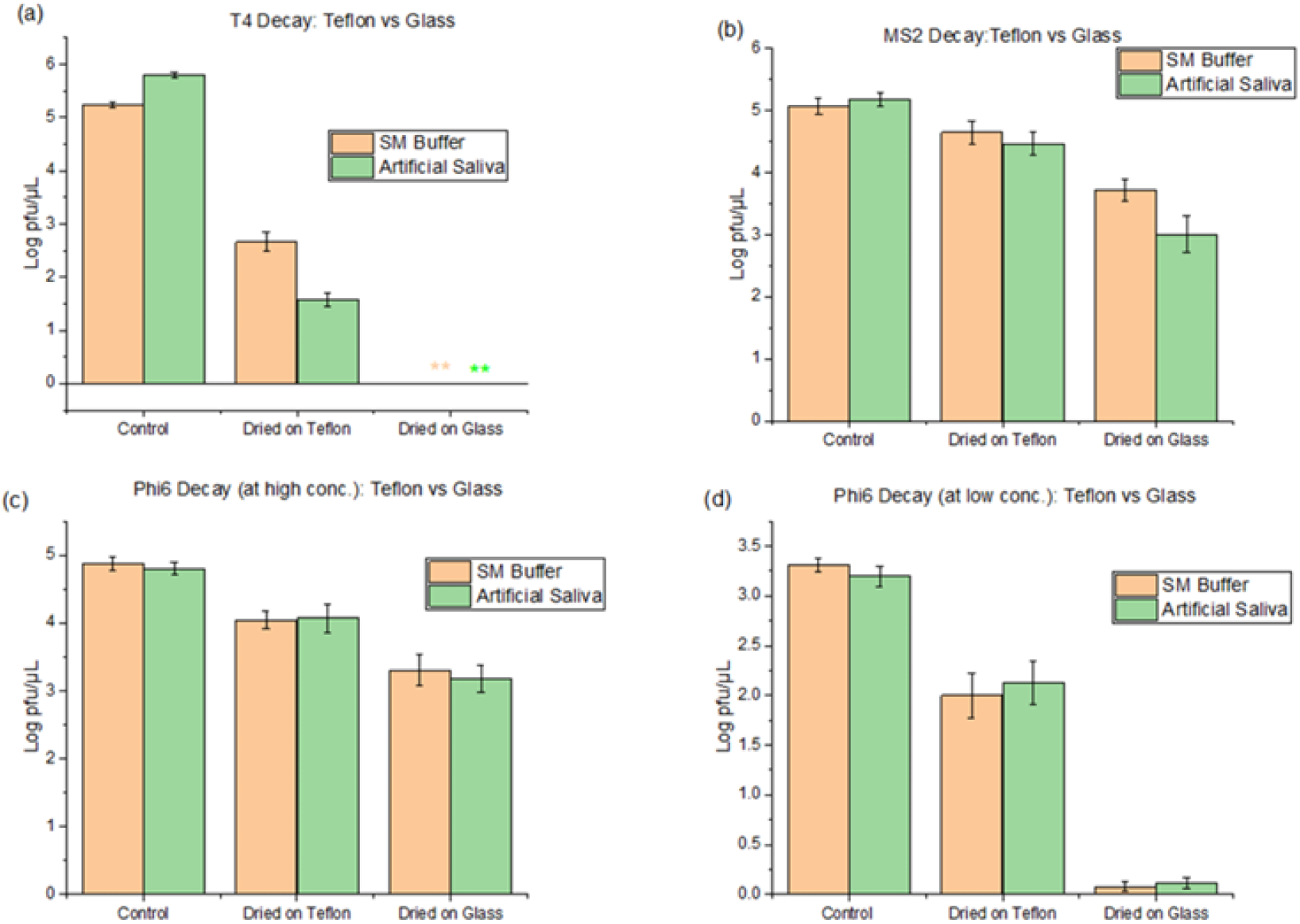
Virus viability on Teflon vs Glass for (a) T4 at ∼ 10^5^ pfu/µL (b) MS2 at ∼ 10^5^ pfu/µL (c) Phi6 at ∼ 10^5^ pfu/µL (c) Phi6 at ∼ 10^3^ pfu/µL

Droplets spreading on glass, as opposed to spherical droplets on Teflon (see Fig. 1a), lead to faster evaporation. At intermediate humidity levels, this increased evaporation results in greater virus decay (28,40). Similar findings have been observed regarding bacterial viability when bacteria-laden surrogate respiratory fluid droplets were dried on glass compared to airborne contact-free droplets (41). In this study, bacteria persisted longer in airborne contact-free droplets than in those deposited on glass. This supports the use of Teflon, which has a hydrophobic surface, as a relevant model for studying contactless droplet drying. The higher virus persistence on Teflon compared to glass suggests that, for droplets of the same volume, airborne droplets can remain infectious for a longer duration than those deposited on surfaces. This increased persistence enhances their potential to cause new infections, as these infectious aerosols may be inhaled by susceptible individuals.

### Effect of initial viral load or concentration

Although the initial viral load (high vs. low) did not significantly affect the relative persistence of Phi6 on Teflon, there was a notable difference on glass (see Fig. 5a). It can also be inferred that the disparity in viral persistence between Teflon and glass was considerable at a low viral load; however, at a higher viral load, their persistence was relatively similar. Revisiting the previous assumption (Teflon: airborne, glass: fomite), it can be concluded that at an initial high viral load, the survival of the virus on fomites can be substantial.

**Figure 5.**
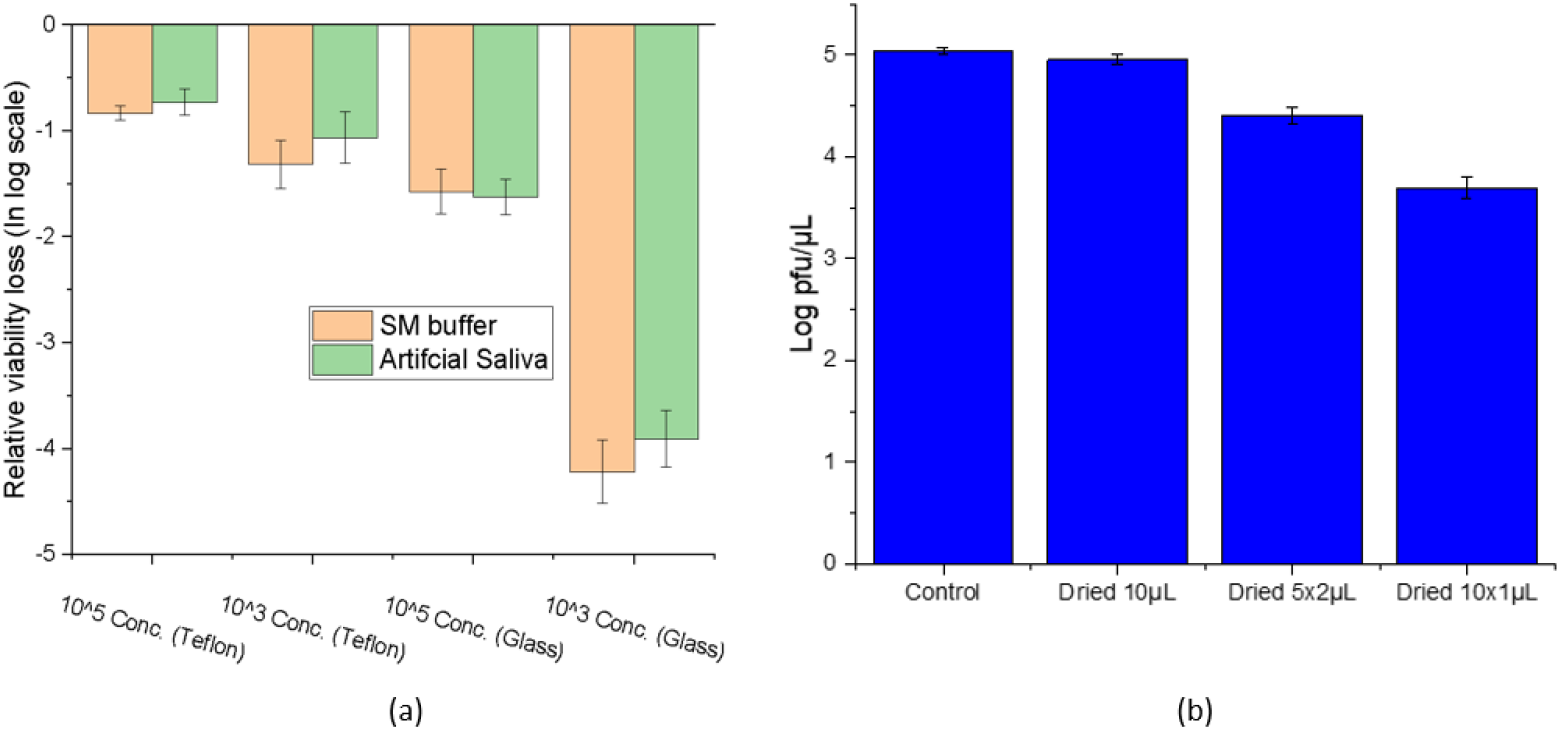
(a) Viral load-dependent viability loss of Phi6 on Teflon and glass, (b) Droplet carrier dependency of Phi6 viability at fixed cumulative droplet volume of 10 µL but different droplet sizes

The primary question is whether the high viral loads observed are an accurate representation of real-world scenarios. Viral load has been identified as a significant factor in virus transmission (42), and the peak in viral load often corresponds with the onset of disease symptoms (43). This stage of the infection cycle carries the highest risk of transmission due to increased virus shedding.

Puhach et al. found substantial variability in viral load among individuals infected with different SARS-CoV-2 strains, ranging from 10^6^ to 10^10^ genome copies per mL (44). Wölfel et al. reported that the average RNA copies for SARS-CoV-2 in sputum samples were approximately 7.00 × 10^6^ copies/mL, with some samples reaching as high as 2.35 × 10^9^ copies/mL (45). Peiris et al. found that the mean viral titer for SARS-CoV ranged from 5.1 × 10^1^ copies/mL to 8.9 × 10^4^ copies/mL over a 23-day period following the onset of illness in patient fecal samples (46).

Tsang et al. reported that the most infectious 20% of cases exhibit 3.1-fold higher infectiousness than average cases (47). Furthermore, Geng and Wang stated that 2% of individuals with SARS-CoV-2 have a viral load exceeding 10^10^/mL, accounting for 90% of the virions circulating within communities (9).

Given the three orders of variability in viral load among infected individuals, fomites originating from those shedding a high viral load still pose a significant risk. The combination of a higher viral load and longer survival time can keep these fomites infectious for extended periods. For example, fomites from ten individuals with average viral loads may be neutralized under normal environmental conditions, but those from a single individual with a high viral load will remain infectious for a longer time. Additionally, a larger volume fraction of respiratory emissions tends to settle down compared to what remains airborne (48, 49), further emphasizing the concern over high viral load fomites. While we do not intend to downplay the importance of airborne transmission, the significantly longer survival of larger respiratory droplets carrying high viral loads could still play a substantial role in the transfer of infection. From a survival perspective, it is essential to focus on the frequency of high viral load individuals or the overall average viral load within an infected cohort in a given region. This information could serve as an indicator of how quickly the disease may spread and what types of interventions may be necessary to mitigate transmission.

### Effect of carrier droplet size

In our study examining droplet size dependency, we observed that the viability of the Phi6 virus was higher in larger droplets compared to smaller ones (10 µL > 5×2 µL > 10×1 µL) when dried on Teflon, as demonstrated in Figure 5(b). The lower survival rate of Phi6 in smaller droplets indicates that virus infectivity is inversely related to the surface area of the carrier droplets. These results further support the notion that the drying rate is the primary environmental factor affecting viability.

This indicates that the drying process leads to a higher loss of infectivity when the surface area to volume ratio of carrier droplets is increased. Applying this to airborne situations, we can conclude the following: For a given total volume, viruses contained in smaller droplets will experience a greater loss of viability due to their faster evaporation compared to those present in larger droplets. Since coughs and sneezes produce droplets with a wide range of diameters, our findings suggest that larger droplets have a higher infectivity factor compared to smaller ones, although we are unable to provide precise quantitative values at this stage.

If this trend holds for droplets smaller than 100 µm, it suggests that these smaller droplets may be less infectious during airborne transmission. However, their prolonged persistence in the air could offset this effect after they completely evaporate, leaving dried residues (Wang et al., 2021) (1). Meanwhile, Anand et al. argued that due to the range of viral load (5 × 10^6– 5 × 10^10 per mL) found in biological fluids, smaller aerosols (less than 5 µm) are unlikely to carry viruses, according to Poisson probability (50). Thus, there may be an optimal size range for droplets that is large enough to carry a viral load capable of surviving the loss of viability due to drying but small enough to evaporate completely before reaching the ground. These droplets would therefore be the most infectious during airborne transmission, considering both virus viability and air persistence.

Droplets in an airborne state generally have a smaller volume (or diameter) compared to those deposited on surfaces, which implies a higher drying rate for the airborne droplets. This higher evaporation rate can reduce the viability of viruses contained in those droplets. Therefore, the overall viability of a mix of droplet sizes (a polydisperse droplet system) faces competing influences: while larger droplets that settle on surfaces spread and remain viable longer, smaller airborne droplets evaporate quickly, reducing the viability of any viruses they carry.

## CONCLUSION

Our findings indicate that for droplets of a given volume, viruses are more likely to remain viable while airborne than when they settle onto surfaces. It is noteworthy that although the viability of viruses decreases with droplet size in the airborne state, overall infectivity may still be considerable. This is due to the fact that smaller droplets can remain suspended in the air for extended periods and can penetrate deeper into the respiratory tract. Furthermore, we observed that at initial viral concentrations of 10^5^/µL, virus survival in both spherical and flat droplets was quite similar, highlighting an increased risk of fomite transmission when the initial viral load is elevated.

## ACKNOWLEDGEMENT

SN acknowledges IIT Bombay for Teaching Assistantship. The project was funded in part by SERB, Department of Science and Technology, India under grant #CRG/2022/004288 (MT) and Department of Biotechnology, India under grant #BT/PR48280/MED/32/829/2023 (MT), and TIH Foundation for IoT and IoE (TIH-IOT), grant # TIH-IoT/2023-3/TDP/HA/SL-IAQ-011 (KK)

To assist the writing process, these Grammarly AI prompts were used: **Prompts created by Grammarly** - *“Improve it”* - *“Make it sound academic”*

## References

1. Wang, C. C., Prather, K. A., Sznitman, J., Jimenez, J. L., Lakdawala, S. S., Tufekci, Z., & Marr, L. C. (2021). Airborne transmission of respiratory viruses. Science, 373(6558),eabd9149

2. Goldman E. (2020). Exaggerated risk of transmission of COVID-19 by fomites. The Lancet. diseases, x3099(20)30561-2 20(8), 892–893. 10.1016/S1473

3. Morawska, L., & Cao, J. (2020). Airborne transmission of SARS-CoV-2: The world should face the reality. Environment international, 139, 105730. 10.1016/j.envint.2020.105730

4. Duval, D., Palmer, J. C., Tudge, I., Pearce-Smith, N., O’Connell, E., Bennett, A., & Clark, R. (2022). Long distance airborne transmission of SARS-CoV-2: rapid systematic review. BMJ (Clinical research ed.), 377, e068743. 10.1136/bmj-2021-068743

5. Zhao, X., Liu, S., Yin, Y., Zhang, T. T., & Chen, Q. (2022). Airborne transmission of COVID-19 virus in enclosed spaces: An overview of research methods. Indoor air, 32(6), e13056. 10.1111/ina.13056

6. Pastorino, B., Touret, F., Gilles, M., de Lamballerie, X., & Charrel, R. N. (2020). Prolonged Infectivity of SARS-CoV-2 in Fomites. Emerging infectious diseases, 26(9), 2256–2257. 10.3201/eid2609.201788

7. Liu, H., Fei, C., Chen, Y., Luo, S., Yang, T., Yang, L., Liu, J., Ji, X., Wu, W., & Song, J. (2021). Investigating SARS-CoV-2 persistent contamination in different indoor environments. Environmental research, 202, 111763. 10.1016/j.envres.2021.111763

8. Hirose, R., Ikegaya, H., Naito, Y., Watanabe, N., Yoshida, T., Bandou, R., Daidoji, T., Itoh, Y., & Nakaya, T. (2021). Survival of Severe Acute Respiratory Syndrome Coronavirus 2 (SARS-CoV-2) and Influenza Virus on Human Skin: Importance of Hand Hygiene in Coronavirus Disease 2019 (COVID-19). Clinical infectious diseases : an official publication of the Infectious Diseases Society of America, 73(11), e4329–e4335. 10.1093/cid/ciaa1517

9. Geng, Y., & Wang, Y. (2023). Stability and transmissibility of SARS-CoV-2 in the environment. Journal of medical virology, 95(1), e28103. 10.1002/jmv.28103

10. Port, J. R., Yinda, C. K., Owusu, I. O., Holbrook, M., Fischer, R., Bushmaker, T., Avanzato, V. A., Schulz, J. E., Martens, C., van Doremalen, N., Clancy, C. S., & Munster, V. J. (2021). SARS-CoV-2 disease severity and transmission efficiency is increased for airborne compared to fomite exposure in Syrian hamsters. Nature communications, 12(1), 4985. 10.1038/s41467-021-25156-8

11. Cheng, P., Luo, K., Xiao, S., Yang, H., Hang, J., Ou, C., Cowling, B. J., Yen, H. L., Hui, D. S., Hu, S., & Li, Y. (2022). Predominant airborne transmission and insignificant fomite transmission of SARS-CoV-2 in a two-bus COVID-19 outbreak originating from the same pre-symptomatic index case. Journal of hazardous materials, 425, 128051. 10.1016/j.jhazmat.2021.128051

12. Greenhalgh, Trisha et al. Ten scientific reasons in support of airborne transmission of SARS-CoV-2 The Lancet, Volume 397, Issue 10285, 1603 – 1605

13. Goldman E. (2021). SARS Wars: the Fomites Strike Back. Applied and environmental microbiology, 87(13), e0065321. 10.1128/AEM.00653-21

14. Firquet, S., Beaujard, S., Lobert, P. E., Sané, F., Caloone, D., Izard, D., & Hober, D. (2015). Survival of Enveloped and Non-Enveloped Viruses on Inanimate Surfaces. Microbes and Environments, 30(2), 140–144

15. Zuo, Z., Kuehn, T. H., Bekele, A. Z., Mor, S. K., Verma, H., Goyal, S. M., Raynor, P. C., & Pui, D. Y. (2014). Survival of airborne MS2 bacteriophage generated from human saliva, artificial saliva, and cell culture medium. Applied and environmental microbiology, 80(9), 2796–2803. 10.1128/AEM.00056-14

16. Lin, K., & Marr, L. C. (2020). Humidity-Dependent Decay of Viruses, but Not Bacteria, in Aerosols and Droplets Follows Disinfection Kinetics. Environmental science & technology, 54(2), 1024–1032. 10.1021/acs.est.9b04959

17. van Doremalen, N., Bushmaker, T., Morris, D. H., Holbrook, M. G., Gamble, A., Williamson, B. N., Tamin, A., Harcourt, J. L., Thornburg, N. J., Gerber, S. I., Lloyd-Smith, J. O., de Wit, E., & Munster, V. J. (2020). Aerosol and surface stability of HCoV-19 (SARS-CoV-2) compared to SARS-CoV-1. medRxiv : the preprint server for health sciences, 2020.03.09.20033217. 10.1101/2020.03.09.20033217

18. Bangiyev, R., Chudaev, M., Schaffner, D. W., & Goldman, E. (2021). Higher Concentrations of Bacterial Enveloped Virus Phi6 Can Protect the Virus from Environmental Decay. Applied and environmental microbiology, 87(21), e0137121. 10.1128/AEM.01371-21

19. Drossinos, Y., Weber, T. P., & Stilianakis, N. I. (2021). Droplets and aerosols: An artificial dichotomy in respiratory virus transmission. Health science reports, 4(2), e275. 10.1002/hsr2.275

20. Anand, S., & Mayya, Y. S. (2020). Size distribution of virus laden droplets from expiratory ejecta of infected subjects. Scientific reports, 10(1), 21174. 10.1038/s41598-020-78110-x

21. Zuo Z.,,Kuehn T., Verma H., Kumar S., Goyal S., Appert J., Raynor P., Ge S., & Pui D. (2013) Association of Airborne Virus Infectivity and Survivability with its Carrier Particle Size, Aerosol Science and Technology, 47:4, 373–382

22. Nayak, S., Mayya, Y., & Tirumkudulu, M. S. (2025). Shrinkage ratios and effective densities of residues formed from drying of simulated expiratory droplets. Journal of Aerosol Science, 184, 106499. 10.1016/j.jaerosci.2024.106499

23. Alonso C, Raynor PC, Davies PR, Torremorell M (2015) Concentration, Size Distribution, and Infectivity of Airborne Particles Carrying Swine Viruses. PLOS ONE 10(8): e0135675. 10.1371/journal.pone.0135675

24. Brown, J. R., Tang, J. W., Pankhurst, L., Klein, N., Gant, V., Lai, K. M., McCauley, J., & Breuer, J. (2015). Influenza virus survival in aerosols and estimates of viable virus loss resulting from aerosolization and air-sampling. The Journal of hospital infection, 91(3), 278–281. 10.1016/j.jhin.2015.08.004

25. Alsved, M., Widell, A., Dahlin, H., Karlson, S., Medstrand, P., & Löndahl, J. (2020). Aerosolization and recovery of viable murine norovirus in an experimental setup. Scientific reports, 10(1), 15941. 10.1038/s41598-020-72932-5

26. Humphrey B, Tezak M, Lobitz M, Hendricks A, Sanchez A, Zenker J, Storch S, Davis RD, Ricken B, Cahill J. Viral Preservation with Protein-Supplemented Nebulizing Media in Aerosols. Appl Environ Microbiol. 2023 Mar 29;89(3):e0154522. doi: 10.1128/aem.01545-22. Epub 2023 Mar 1. PMID: 36856430; PMCID: PMC10057872.

27. Baker, C. A., Gutierrez, A., & Gibson, K. E. (2022). Factors Impacting Persistence of Phi6 Bacteriophage, an Enveloped Virus Surrogate, on Fomite Surfaces. Applied and environmental microbiology, 88(7), e0255221. 10.1128/aem.02552-21

28. Cunliffe, A. J., Wang, R., Redfern, J., Verran, J., & Ian Wilson, D. (2022). Effect of environmental factors on the kinetics of evaporation of droplets containing bacteria or viruses on different surfaces. Journal of Food Engineering, 336, 111195. 10.1016/j.jfoodeng.2022.111195

29. Schaub, A., David, S. C., Glas, I., Klein, L. K., Violaki, K., Terrettaz, C., Motos, G., Bluvshtein, N., Luo, B., Pohl, M., Hugentobler, W., Nenes, A., Krieger, U. K., Peter, T., Stertz, S., & Kohn, T. (2024). Impact of organic compounds on the stability of influenza A virus in deposited 1-μL droplets. mSphere, 9(9), e0041424. 10.1128/msphere.00414-24

30. Fedorenko, A., Grinberg, M., Orevi, T. et al. Survival of the enveloped bacteriophage Phi6 (a surrogate for SARS-CoV-2) in evaporated saliva microdroplets deposited on glass surfaces. Sci Rep 10, 22419 (2020). 10.1038/s41598-020-79625-z

31. Monika, Damle, E. A., Kondabagil, K., & Kunwar, A. (2025). Comparative study of inactivation efficacy of far-UVC (222 nm) and germicidal UVC (254 nm) radiation against virus-laden aerosols of artificial human saliva. Photochemistry and photobiology, 10.1111/php.14062. Advance online publication. 10.1111/php.14062

32. Gardner, A., Ghosh, S., Dunowska, M., & Brightwell, G. (2021). Virucidal Efficacy of Blue LED and Far-UVC Light Disinfection against Feline Infectious Peritonitis Virus as a Model for SARS-CoV-2. Viruses, 13(8), 1436. 10.3390/v13081436

33. Cox CS, Harris WJ, Lee J. 1974. Viability and electron microscope studies of phages T3 and T7 subjected to freeze-drying, freeze-thawing and aerosolization. J. Gen. Microbiol. 81:207–215.

34. Tzlil, S., Kindt, J. T., Gelbart, W. M., & Ben-Shaul, A. (2003). Forces and pressures in DNA packaging and release from viral capsids. Biophysical journal, 84(3), 1616–1627. 10.1016/S0006-3495(03)74971-6

35. Harper GJ. 1963. The influence of environment on the survival of airborne virus particles in the laboratory. Arch. Virol. 13:64 –71.

36. Benbough JE. 1971. Some factors affecting the survival of airborne viruses. J. Gen. Virol. 10:209 –220

37. Kormuth, K. A., Lin, K., Prussin, A. J., Vejerano, E. P., Tiwari, A. J., Cox, S. S.,Myerburg, M. M., Lakdawala, S. S., & Marr, L. C. (2018). Influenza Virus Infectivity is retained in Aerosols and Droplets Independent of Relative Humidity. The Journal of infectious diseases, 218(5), 739–747. 10.1093/infdis/jiy221

38. Cordomi A, Edholm O, Perez JJ. 2008. Effect of ions on a dipalmitoyl phosphatidylcholine bilayer. A molecular dynamics simulation study. J.Phys. Chem. B 112:1397–1408.

39. Lee SJ, Song Y, Baker NA. 2008. Molecular dynamics simulations of asymmetric NaCl and KCl solutions separated by phosphatidylcholine bilayers: potential drops and structural changes induced by strong Na+lipid interactions and finite size effects. Biophys. J. 94:3565–3576

40. French, A. J., Longest, A. K., Pan, J., Vikesland, P. J., Duggal, N. K., Marr, L. C., & Lakdawala, S. S. (2023). Environmental Stability of Enveloped Viruses Is Impacted by Initial Volume and Evaporation Kinetics of Droplets. mBio, 14(2), e0345222. 10.1128/mbio.03452-22

41. Amey Nitin Agharkar, Dipasree Hajra, Durbar Roy, Vivek Jaiswal, Prasenjit Kabi, Dipshikha Chakravortty, Saptarshi Basu (2024). Evaporation of bacteria-laden surrogate respiratory fluid droplets: On a hydrophilic substrate vs contact-free environment confers differential bacterial infectivity. Physics of Fluids 1 March; 36 (3): 031912. 10.1063/5.0196219

42. Marc, A., Kerioui, M., Blanquart, F., Bertrand, J., Mitjà, O., Corbacho-Monné, M., Marks, M., & Guedj, J. (2021). Quantifying the relationship between SARS-CoV-2 viral load and infectiousness. eLife, 10, e69302. 10.7554/eLife.69302

43. Puhach, O., Meyer, B., & Eckerle, I. (2023). SARS-CoV-2 viral load and shedding kinetics. Nature reviews. 10.1038/s41579-022-00822-w Microbiology, 21(3), 147–161. 16

44. Puhach, O., Adea, K., Hulo, N., Sattonnet, P., Genecand, C., Iten, A., Jacquérioz, F., Kaiser, L., Vetter, P., Eckerle, I., & Meyer, B. (2022). Infectious viral load in unvaccinated and vaccinated individuals infected with ancestral, Delta or Omicron SARS-CoV-2. Nature medicine, 28(7), 1491–1500. 10.1038/s41591022-01816-0

45. Wölfel, R., Corman, V. M., Guggemos, W., Seilmaier, M., Zange, S., Müller, M. A., Niemeyer, D., Jones, T. C., Vollmar, P., Rothe, C., Hoelscher, M., Bleicker, T., Brünink, S., Schneider, J., Ehmann, R., Zwirglmaier, K., Drosten, C., & Wendtner, C. (2020). Virological assessment of hospitalized patients with COVID 2019. Nature, 581(7809), 465–469. 10.1038/s41586-020-2196-x

46. Peiris, J. S., Chu, C. M., Cheng, V. C., Chan, K. S., Hung, I. F., Poon, L. L., Law, K. I., Tang, B. S., Hon, T. Y., Chan, C. S., Chan, K. H., Ng, J. S., Zheng, B. J., Ng, W. L., Lai, R. W., Guan, Y., Yuen, K. Y., & HKU/UCH SARS Study Group (2003). Clinical progression and viral load in a community outbreak of coronavirus-associated SARS pneumonia: a prospective study. Lancet (London, England), 361(9371), 1767–1772. 10.1016/s0140-6736(03)13412-5

47. Tsang, T. K., Huang, X., Wang, C., Chen, S., Yang, B., Cauchemez, S., & Cowling, B. J. (2023). The effect of variation of individual infectiousness on SARS-CoV-2 transmission in households. eLife, 12, e82611. 10.7554/eLife.82611

48. Katre, P., Banerjee, S., Balusamy, S., & Sahu, K. C. (2021). Fluid dynamics of respiratory droplets in the context of COVID-19: Airborne and surfaceborne transmissions. Physics of fluids (Woodbury, N.Y. : 1994), 33(8), 081302. 10.1063/5.0063475

49. Pöhlker, M. L., Krüger, O. O., Förster, J.-D., Berkemeier, T., Elbert, W., Fröhlich-Nowoisky, J., Pöschl, U., & Pöhlker, C. (2023). Respiratory aerosols and droplets in the transmission of infectious diseases. Reviews of Modern Physics, 95(4), 045001. 10.1103/RevModPhys.95.045001

50. Anand, S., Krishan, J., Sreekanth, B., & Mayya, Y. S. (2022). A comprehensive modelling approach to estimate the transmissibility of coronavirus and its variants from infected subjects in indoor environments. Scientific reports, 12(1), 14164. 10.1038/s41598-022-17693-z

